# Monitoring the depth of anesthesia using Autoregressive model and Sample entropy

**DOI:** 10.1101/634675

**Authors:** Fu G. Zhu, Xiao G. Luo, Chang J. Hou, Dan Q. Huo, Peng Dang

**Affiliations:** Key Laboratory of Biorheological Science and Technology, Ministry of Education, Bioengineering college, Chongqing University, Chongqing 400044, China; Chongqing Engineering Research Center of Medical Electronics Technology, Chongqing 400044, China

**Keywords:** electroencephalogram, sample entropy, autoregressive model, weighted k-nearest neighbor, depth of anesthesia

## Abstract

Anesthesia is an important part in modern surgery, and the way how to effectively monitor the depth of anesthesia (DOA) is core issue in the anesthesia work. Since anesthetics mainly affected the brain of patients, it is very effective to monitor DOA by electroencephalogram (EEG). This paper proposes a method for monitoring DOA using EEG. First, the sample entropy (SampEn) of EEG were calculated as a feature vector. Simultaneously, the Burg recursive algorithm was used to solve the autoregressive model (AR model) and AR coefficients were extracted as feature vectors. Later, according to the characteristics of uneven distribution of sample points, the weighted k-nearest neighbor (WKNN) classifier was selected. The Anesthesia was divided into awake, mild, moderate and deep by WKNN classifier. According to the results, the correlation coefficient between the SampEn of the EEG and Bispectral Index (BIS) is above 0.8. There is a correlation between the first five orders of AR coefficient and the BIS index, and the correlation of the second order reaches 0.8. Through the validation of 30 patients, this method can assessment of DOA effectively and accurately.

## Introduction

Considering the safety and efficiency of surgery, it is very necessary to make the research on non-invasive anesthesia depth monitoring. Excessive doses of anesthetic during anesthesia can lead to delayed recovery and coma [1] and insufficient doses of medication may keep the patient status at near consciousness [2]. Thus, guiding the anesthesiologist to use anesthetic agents accurately can rely on the accurate judgment of the anesthesia depth. At the same time, it can ensure the comfort and safety of patients during the surgery [3].

The traditional method of monitoring depth of anesthesia (DOA) is to use clinical features such as breathing, sweating, heart rate, limb movements, blood pressure, and pulse [4]. However, the variation could be generated in these parameters depending on the patient and the surgery type. In addition, clinical features are easily altered with the use of vasodilators and muscle relaxants [5]. Therefore, these clinical features have significant limitations in evaluating DOA.

In the past decades, using electroencephalogram (EEG) signals to monitoring DOA has become a research hotspot. Hutt and Longtin et al [6] found that brain is directly affected by anesthetic drugs, and they proposed the method that using EEG signals to monitor DOA was very efficient, accurate, safe, and non-invasive. Recently, common equipment based on EEG to monitoring DOA mainly contains Bispectral Index (BIS) [7] and M-Entropy module [8] BIS uses a complex algorithm to divide the depth of anesthesia into 0 to 100 index, which was commonly used in hospitals. However, BIS also has some limitations, such as the sensitivity to artifacts [9] and a delay in response to change in EEG [10]. M-Entropy module, developed by Datex-Ohmeda, this method was implemented by calculating the spectral entropy of the EEG signals, and the fast Fourier transformation (FFT) was used in calculating the spectral entropy. Despite FFT is a linear algorithm, many studies have showed that EEG signals are non-linear random signals.

Recently, methods for EEG signal analysis have developed very rapidly, and these methods have been widely used in DOA. Since the EEG signal is a nonlinear signal, the nonlinear method has been favored by researchers, such as Lyapunov exponent [11], L-Z complexity analysis [12], Bayesian method [13,14], Hilbert–Huang transform [15], recurrence analysis [16], Detrended Fluctuation Analysis (DFA) [17], and Entropy [18–20]. Although these methods can initially extract the characteristics of EEG signals, they have a common flaw: none of these parameters can independently monitor DOA. Therefore, many studies extracted the characteristics of EEG signals using multiple parameters [21,22].

This study aim to monitoring DOA using EEG signals, which mainly includes two steps of feature extraction and anesthesia staging. First, AR coefficient and sample entropy were extracted from EEG signals. Then, we classify the EEG signals according to the extracted feature values into different anesthesia states using the WKNN classifier. This method divides the anesthesia status into four different levels like most current research.

## Methods

### Data collection

In this experiment, BIS monitor is used to acquire two channels EEG signals, which produced by Aspect and sampling frequency is 256 Hz. The data was collected via BIS monitor’s matching electrode. All EEG signals include complete data between awake and anesthesia. The BIS index value was recorded to provide clinical proof, while collecting the EEG signals. Thirty patients were collected data, whose ages from twenty to sixty. The data collected by BIS monitor includes EEG signals, EMG signals, location of the EEG when BIS index updated and signal quality information. The patient did not have any brain disease and did not take any drugs that affected the anesthesia before the operation. All patients are aware of the experiment purpose and method of the trial and signed the informed consent form.

### Signal preprocessing

Factors such as environment during the signal acquisition, acquisition method and surgical operation make the collected EEG signals contain a lot of noise. Because the EEG signal is weak and the external noise is generally relatively strong, the features contained in the EEG signal are easily covered by external interference. If noise can not be eliminated, feature vectors will be inaccurate. Empirical Mode Decomposition (EMD) was used when noise is removed. Comparing the original signal and the filtered signal, the baseline drift in the EEG signal can be eliminated.

**Fig 1.**
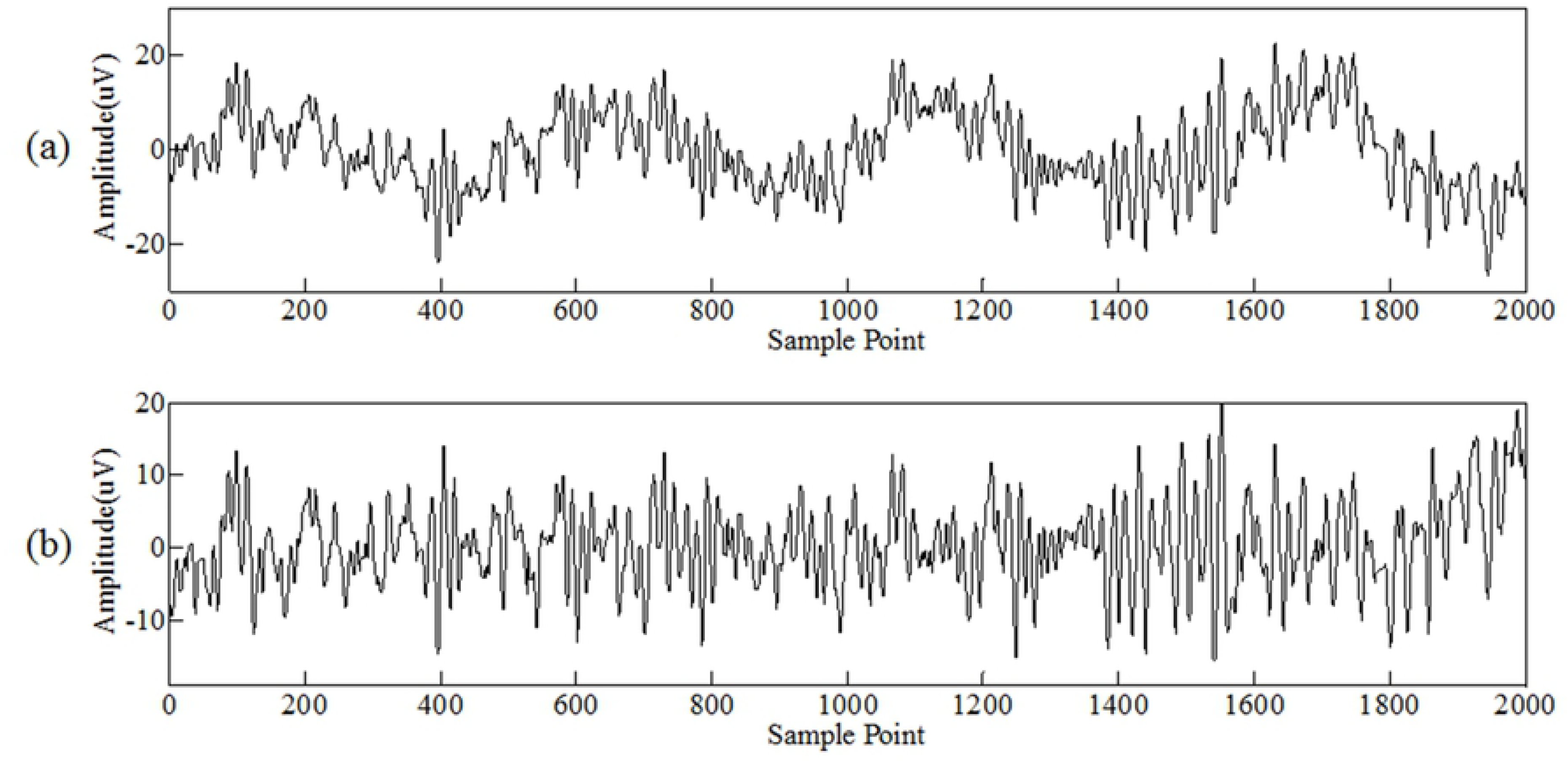
Filtering of noise in EEG signals. (a) Original EEG signals collected during anesthesia. (b) Original EEG signal are filtered by EMD.

EEG signal is segmented by sliding window, the window length is 2560 (10 seconds of EEG signals) and step size is 512 (2 seconds of EEG signals). Then, each segment of EEG signals was analyzed and the feature vectors related to the depth of anesthesia is extracted. In this paper, sample entropy and AR model were used to analyze EEG signals.

**Fig 2.**
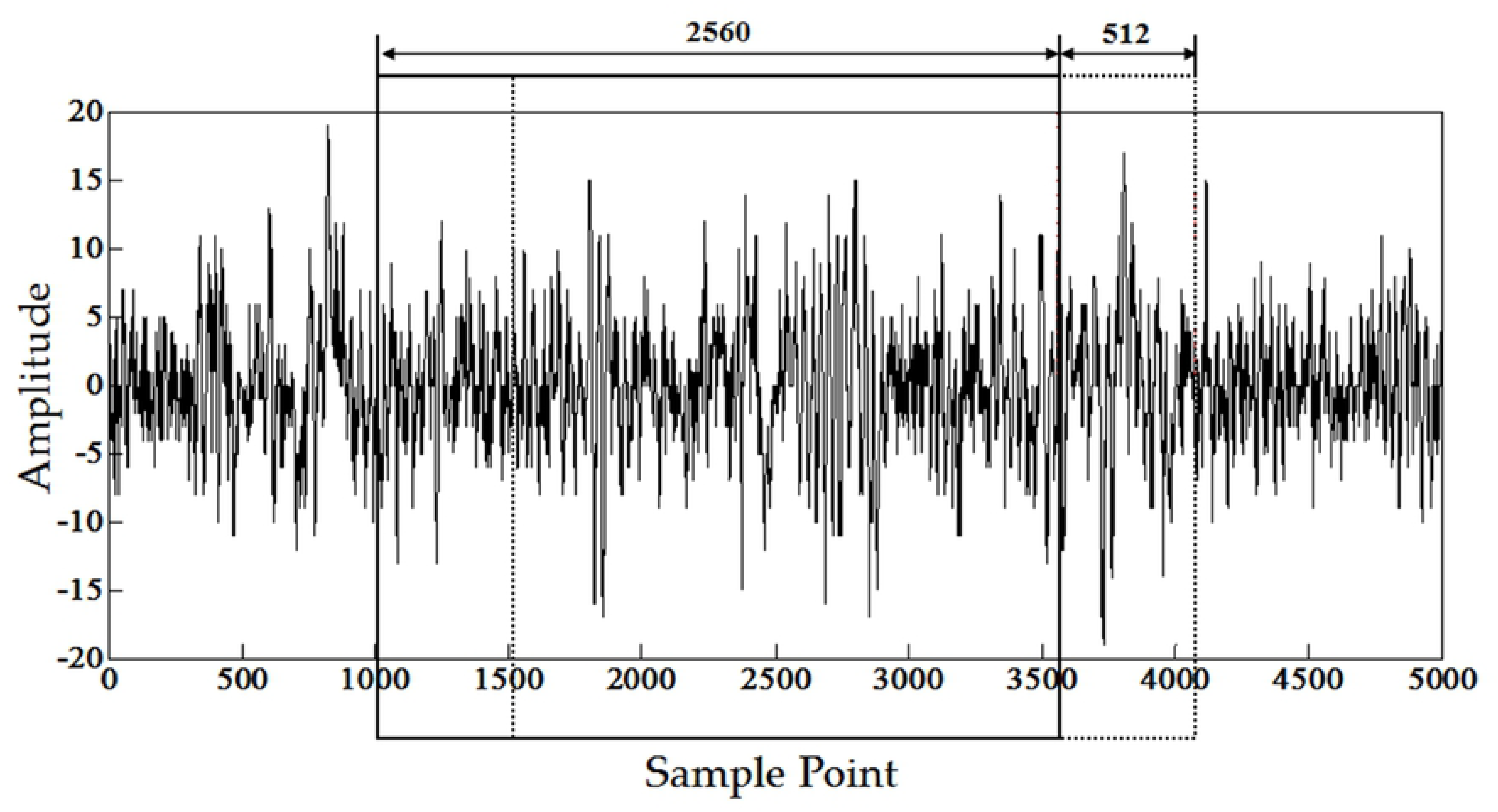
Move window length and step size. The end of the window corresponds to the position of EEG signal when BIS is updated

### Sample entropy

Sample entropy is a nonlinear analysis method proposed by Richman JS and Moorman JR [23], which is a method based on approximate entropy. Sample entropy not only has all the advantages of approximate entropy, but also avoids statistical inconsistencies in approximate entropy. Due to the non-nonlinear characteristic of EEG data, sample entropy is widely used in the analysis of EEG signals [24]. Given the one-dimensional time series. {*x(i)*} *i=1,2,…,N*, the algorithm for sample entropy is as follows:

1. Extracting *N-m*+*1* vectors *Y*_*m*_(*i*) from *X(i)* is defined as:

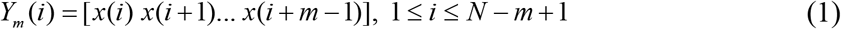
2. Compute the distance between Y_m_(i) and Y_m_(j) for each i defined as *d*[*Y*_*m*_(*i*), *Y*_*m*_(*j*)]

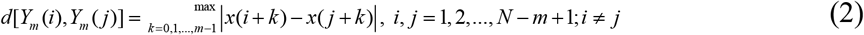
3. Given the threshold *r* (*r>0*), Calculate the number of *d*[*Y*_*m*_(*i*),*Y*_*m*_(*j*)] less than *r* as 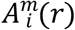, and the 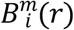 is defined as:

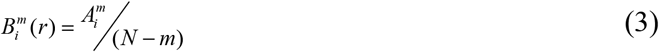
4. Calculate the average value of 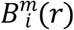:

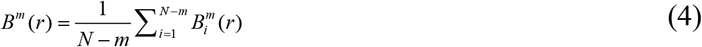
5. Let *m*=*m*+*1*, repeat 1)~4), get *B*^*m* + 1^(*r*)
6. The sample entropy of the *x(i)* is:

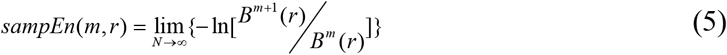

In the actual calculation, since the sequence length is limited, the estimated sample entropy of the sequence is obtained:

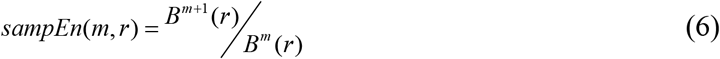

Obviously, the value of the sample entropy is related to m and r. Usually take *m=2* or *1*, *r=0.1~0.25 sd* (*sd* is the standard deviation of the sequence)[25], In this study, the value of *m* is *2*, *r* equals 0.2*sd* and signal length *N* is *2560*. The specific data processing process is as follows

First, a sliding window was added to the EEG, which window length is 2560 sample points and window step size is 512 sample points (number of samples taken in 2second). Then, SampEn of the signal in the window was calculated. Final, correlation analysis and trend comparison were conducted according to sample entropy and BIS values.

### Autoregressive model

As an effective method to represent the internal information and characteristics of the signal [26], autoregressive (AR) model plays important role in EEG signals analysis. Such as Brain Computer Interface (BCI) [27], Judgement of schizophrenia [28], hypnosis levels [29] and analysis of sleep [30], epilepsy diagnosis [31]^**Error! Reference source not found.**^ and emotion recognition [32]^**Error! Reference source not found.**^. In AR model each sample has a relationship with all the samples in front of it. The AR model can be expressed as:

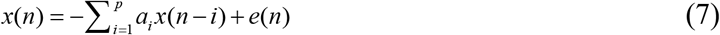

Where: *e(n)* is a white noise; parameter *p* is the order of the AR model and a_i_ is AR coefficient. AR coefficients can reflect the dependency weight of the signal *x(n)* and the previous signal effectively. There are many ways to obtain the AR coefficient. Here, the Burg algorithm is used to obtain the AR coefficient. By analyzing the AR coefficient, five AR coefficients, which have the highest correlation with the BIS value, were used as the characteristics of the EEG signals.

### Weighted K-Nearest Neighbor

K-Nearest Neighbor (KNN) has the advantages of simple algorithm and accurate classification, and is used in many aspects. Its principle is that the category to which the sample belongs determined by its nearest k neighbors. The input sample was classified into the class which is the majority of the k neighbors belongs to. First, the method calculated the Euclidean distance from input point to the points in the data set [33]^**Error! Reference source not found.**^. The expression is as follows:

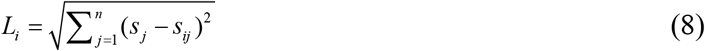

S_j_ is the input point, and S_ij_ is the point in the data set. Then, sort the points in the training set in descending order by distance, and select the first *K* points. Finally, input samples are assigned to a class that has the largest share of *K* points. So the choice of *K* value is extremely critical in KNN algorithm. If the *K* value is too small, insufficient number of adjacent parameter points, and the express feature of the reference point can not be expressed sufficiently which will bring huge errors. If the *K* value is too large, it is easy to select a reference point that is far from the sample point, which leads to inaccurate results.

Based on the KNN, a new algorithm WKNN has been widely used [34,35], and the classification effect of WKNN was better than KNN. Due to the imbalance of sample distributions, the majority of the *K* neighbors had a large sample size when a new sample is input. This phenomenon can lead to misclassification. In this study, the distribution of anesthesia samples at various anesthesia stages is uneven, so WKNN was selected as the classifier.

## Results and Discussions

### Relationship between sample entropy and depth of anesthesia

A section of EEG signals was extracted through the sliding window, and then the SampEn of EEG signals was calculated. To determine whether sample entropy can evaluate the depth of anesthesia, the obtained sample entropy and BIS value trend were used for comparison. As shown in Fig.3a, the BIS value curve is the depth of anesthesia detected by BIS equipment, and the SampEn curve is obtained by linear multiplication. It can be seen from the curves that the trend of sample entropy is consistent with the trend of DOA.

**Fig 3.**
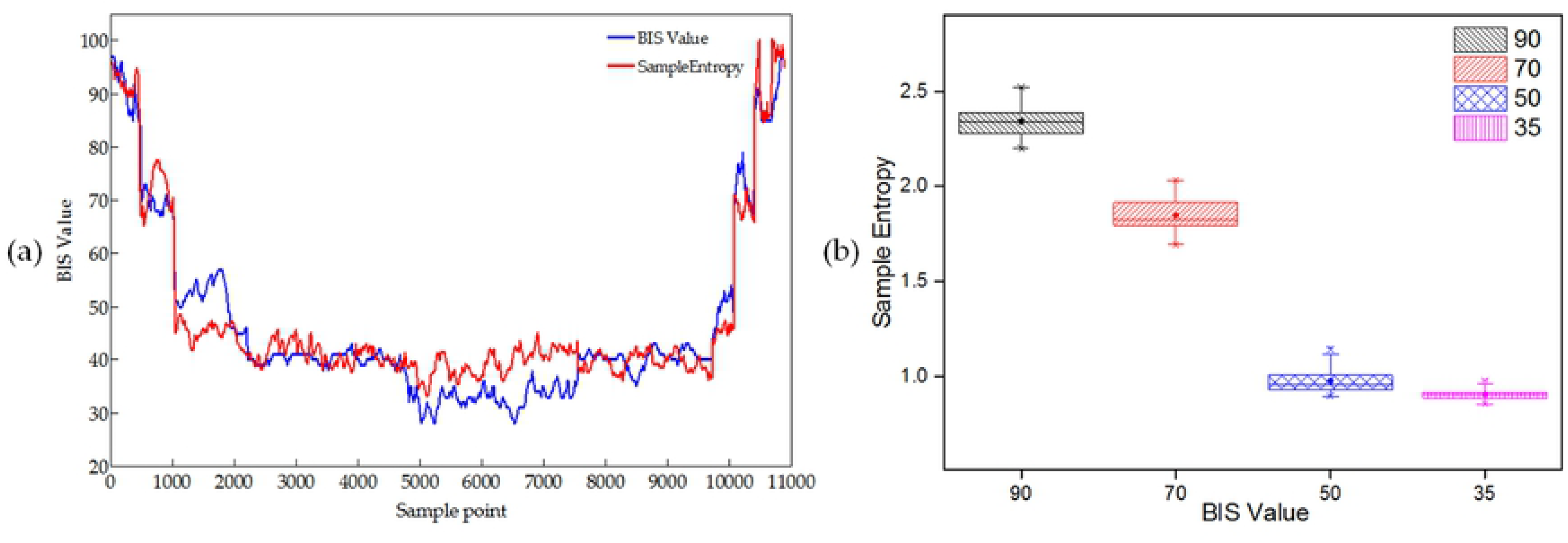
Comparison of results between BIS and Sample Entropy. (a) Sample Entropy and BIS trend. (b) Box chart of BIS value and sample entropy.

To reflect the SampEn of EEG signals at four DOA states directly, SampEn of the EEG is calculated and the values of the four states are about 35, 50, 70 and 90. Fig.3b indicates that the difference between sample entropy of EEG signals during wake, mild and moderate anesthesia is significant. Although sample entropy is overlapped between moderate and deep anesthesia, sample entropy during moderate anesthesia is significantly higher than deep anesthesia.

The correlation between EEG sample entropy and BIS value was analyzed to verify the relationship between EEG signals entropy and DOA. The results show that the correlation is 0.84 and correlation is significant at 0.01 level (2-tiailed). It proves that sample entropy has a strong correlation with BIS value, so it is feasible that use sample entropy to monitor DOA.

### Relationship between sample entropy and depth of anesthesia

The correlation between the AR coefficient and the BIS value was calculated. By comparing the correlations, the AR coefficient associated with BIS was selected to monitor DOA. Table.1 denotes that there is a correlation between the first five orders of AR coefficients and the BIS values and the other AR coefficient has no relationship with the BIS value. Therefore, the AR coefficient of the first five orders is used as the feature vectors for monitoring DOA.

**Table 1.** Correlation between AR coefficient and BIS value

| AR coefficient | 1 | 2 | 3 | 4 | 5 | 6 | 7 | 8 |
| --- | --- | --- | --- | --- | --- | --- | --- | --- |
| Pearson correlation | -.367** | .805** | -.645** | -.594** | -.350** | .129** | 0.056 | 0.056 |
| Sig (2-tailed) | 0 | 0 | 0 | 0 | 0 | 0.001 | 0.159 | 0.162 |
| N | 6670 | 6670 | 6670 | 6670 | 6670 | 6670 | 6670 | 6670 |

\*\*Correlation is significant at 0.01 level (2-tailed)

To reflect the AR coefficient of EEG at four DOA states directly, EEG with BIS values of 35, 50, 70 and 90 were obtained. First five orders of AR coefficients of EEG under different DOA were calculated. Fig.4b indicates that the second order AR coefficient decreased obviously with the increase of anesthesia depth. It can be seen from Fig.4 that another AR coefficients were different under different anesthesia, but the phenomenon was not as obvious as the second order AR coefficient.

**Fig 4.**
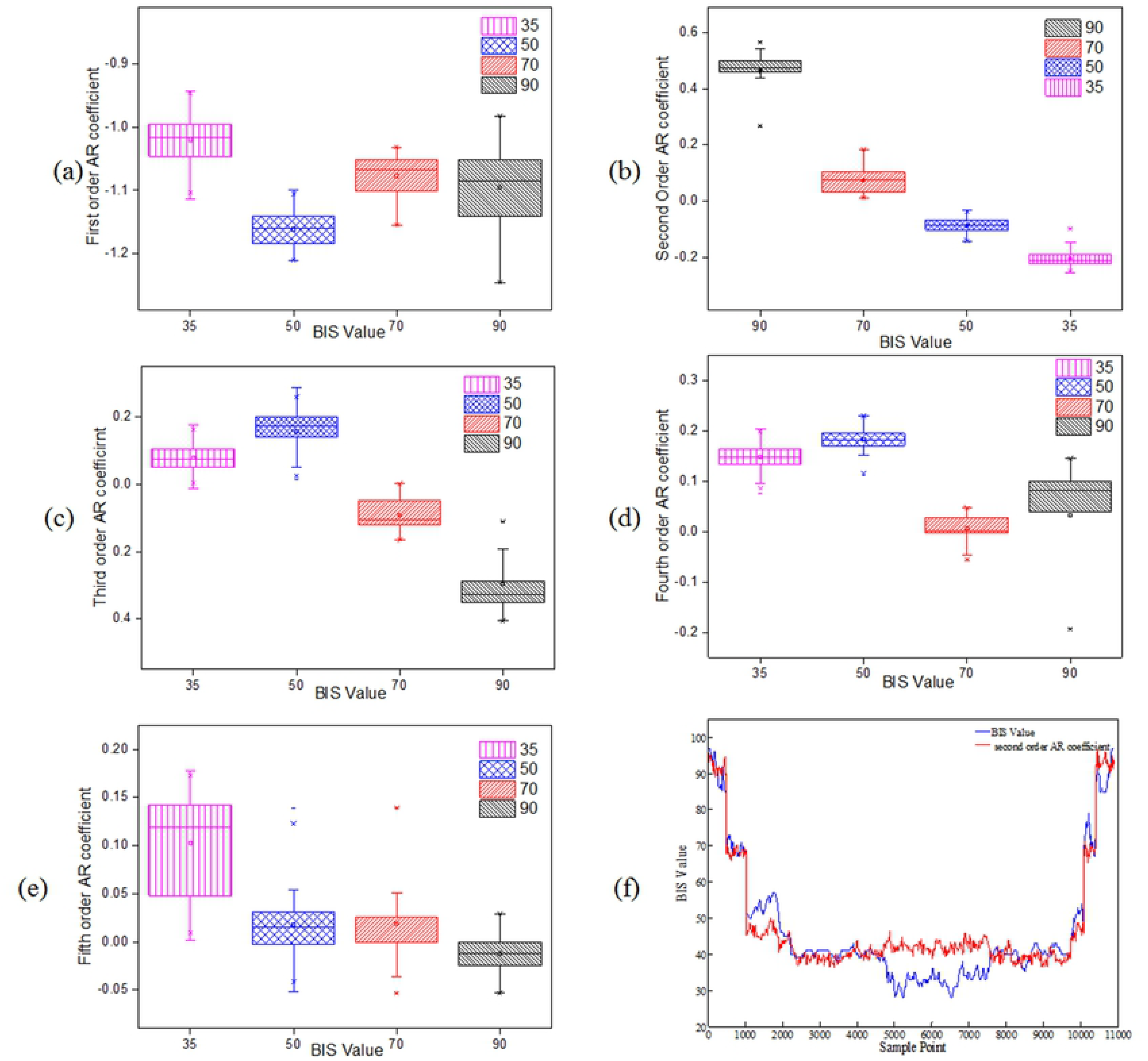
AR coefficient of EEG at four DOA states. (a)~ (e) Box chart of BIS value and first order to fifth order of AR coefficients. (f) Second order of the AR coefficient and BIS trend.

The second order of AR coefficient with the highest correlation was selected to compared with BIS value. In Fig.4f, the second order of AR coefficient curve was obtained by linear multiplication. Fig.4f reflects that, the trend of the AR coefficient and DOA is similar just like the sample entropy. Although there is a difference between the two curves during deep anesthesia, the trend of the curves is consistent with the BIS values during other anesthesia depths. Like sample entropy, the second order of AR coefficients are calculated when anesthesia depth around 35, 50, 70 and 90. The second order of AR coefficients in awake, mild anesthesia, moderate anesthesia, and deep anesthesia are significantly different. Fig.4b illustrates that the second order AR coefficient decreased obviously with the increase of anesthesia depth.

### Feature vector selection and classification

AR coefficient and SampEn were used as feature vectors to select the value of parameter K in WKNN through optimization. The recognition rate shows a trend of increasing first and then decreasing when the K value increases. The recognition rate reaches the maximum when K is equal to seven. WKNN needs to calculate the distance from the test point to all sample points. In order to reduce the calculation amount and improve the recognition rate, the AR coefficient was selected as the perfect combination. As can be seen from Fig.5b, when a single AR coefficient was taken as the feature vector, the recognition rate of the second order of AR coefficient reaches about 75% at the maximum and the recognition rate of the first order of AR coefficient is only about 45% at the minimum. The AR coefficients were combined according to the recognition rate from high to low. The AR coefficient combination, whose recognition rate is optimal, is the second order to the fifth order.

**Fig 5.**
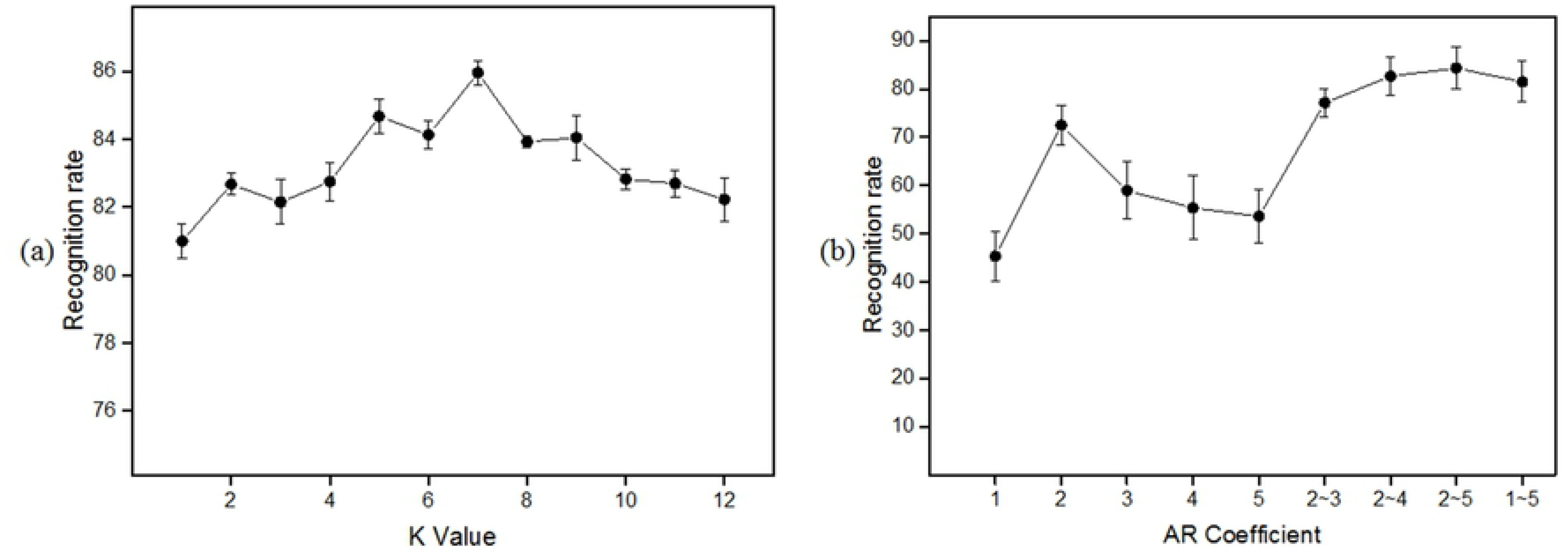
Parameter optimization. (a) K value and recognition rate. (b) AR combination and recognition rate. Error bars represent the standard deviations of the independent measurements

Through applying the AR coefficients and sample entropy as feature vectors, the depth of anesthesia was divided into awake (81-100), mild anesthesia (61-80), moderate anesthesia (41-60), and deep anesthesia (0-40). From the results of Fig.5, the K value is seven in the WKNN and AR coefficients is taken from the second order to the fifth order. In Fig.6, all depths of anesthesia present a high recognition rate. The results show that the recognition rate under deep anesthesia is the lowest, recognition rate under light anesthesia is the highest and the overall recognition rate reaches 88.71%.

**Fig 6.**
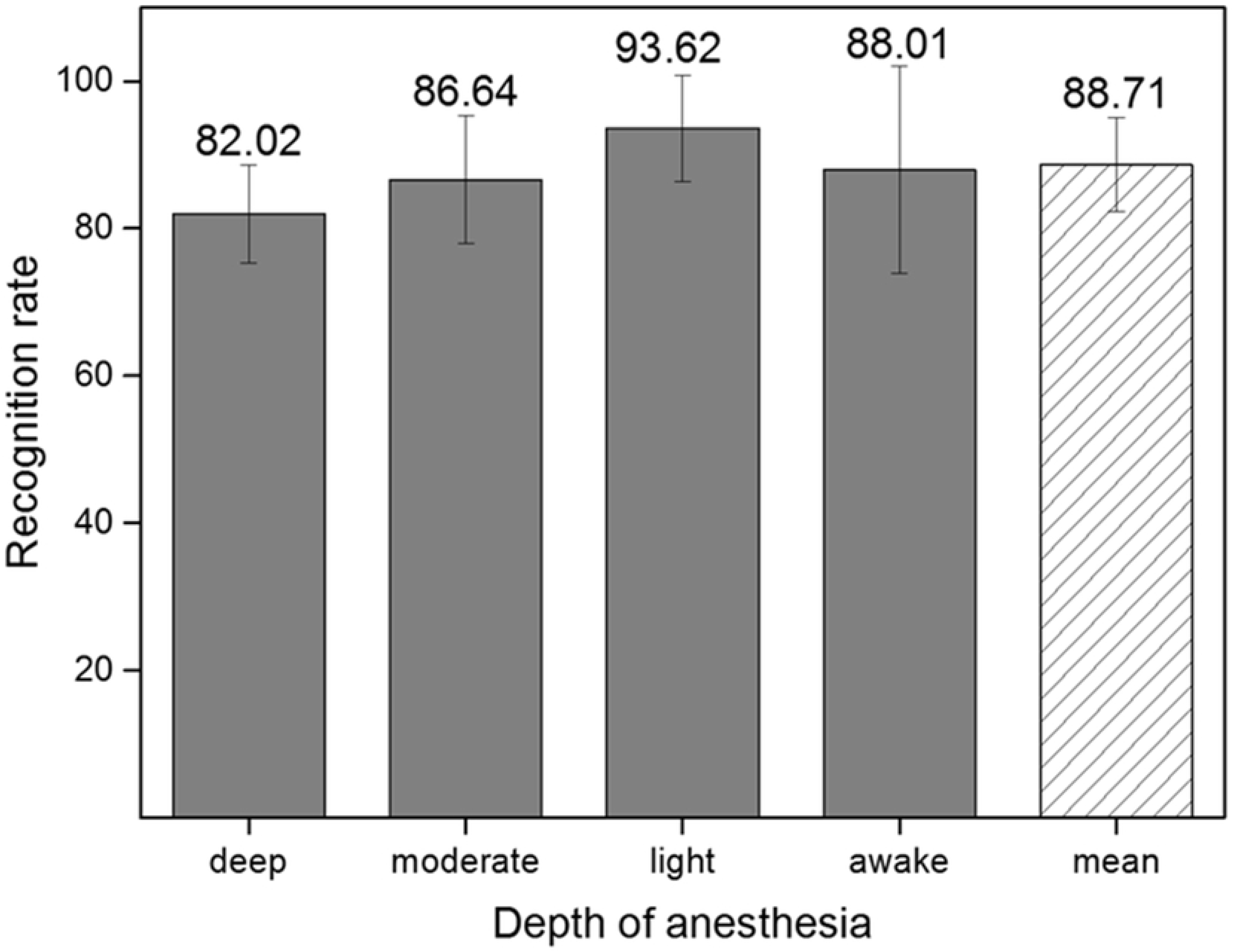
Recognition rate of DOA. WKNNN is used for classification, the feature data constructs a training set and a test set separately according to 4:1, and the recognition rate of each state is counted.

The WKNN classification results of one patient were counted, in order to analyze the distribution of classification results under different anesthesia status. As shown in Fig.7, the same patient has a higher recognition rate for different depth of anesthesia and most of the misidentified samples were identified as being in adjacent categories, which is consistent with the physiological mechanisms of EEG signals. In the four anesthesia status, the recognition rate in awake status is poor which more than 20% samples were mistaken for light anesthesia, and the recognition rate under light anesthesia is best which only very few samples were identified incorrectly.

**Fig 7.**
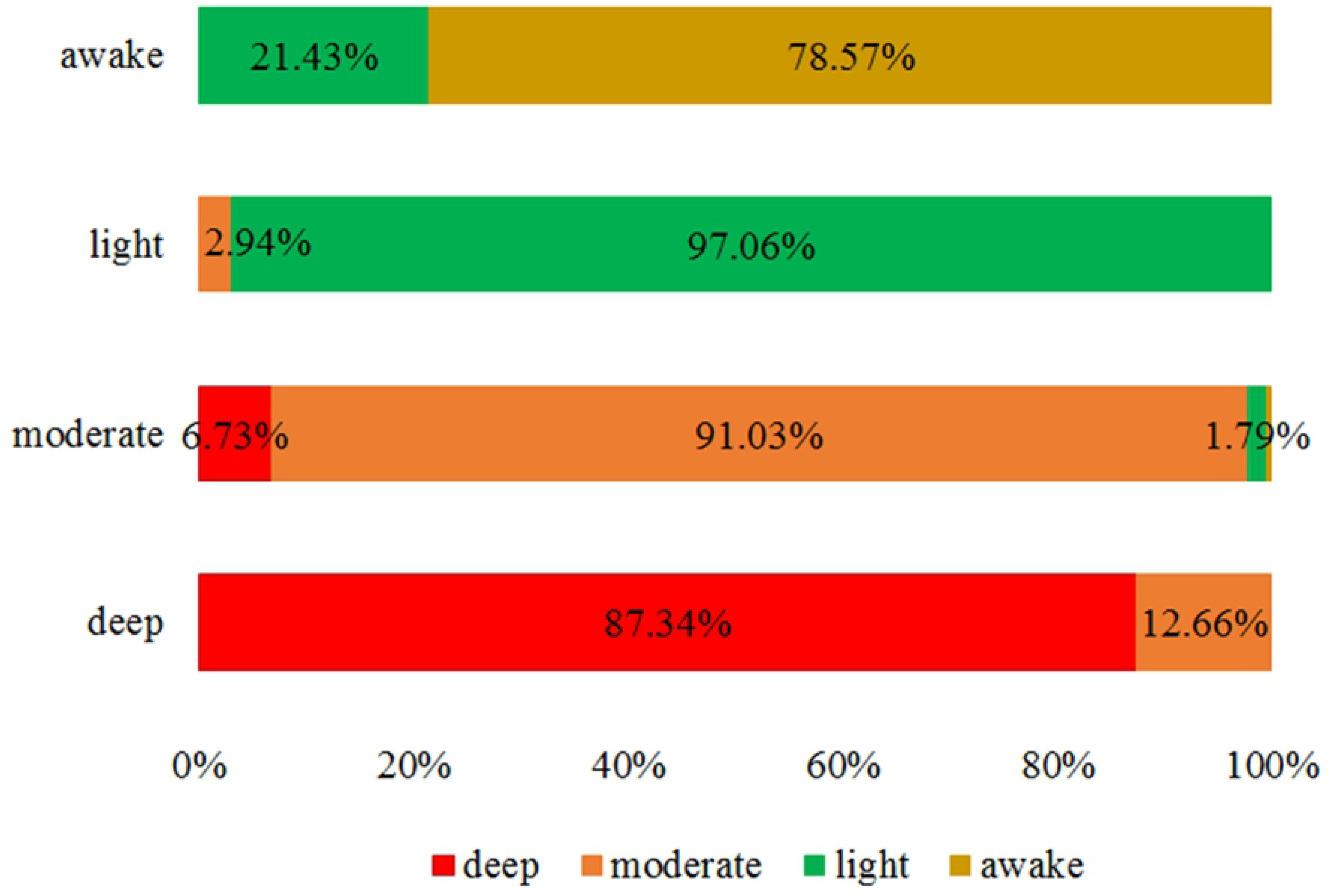
WKNN classification results of one patient under different DOA. The sample lengths were different under anesthesia four kinds of DOA.

In order to observe the spatial distribution of eigenvalues at different depths of anesthesia visually, the spatial distribution scatter plot was drawn using two feature vectors (Sample entropy and second order AR coefficient) which have the highest similarity with the BIS index. In Fig.8, although the feature vectors extracted under the four anesthesia states have cross distribution, they show obvious clusters.

**Fig 8.**
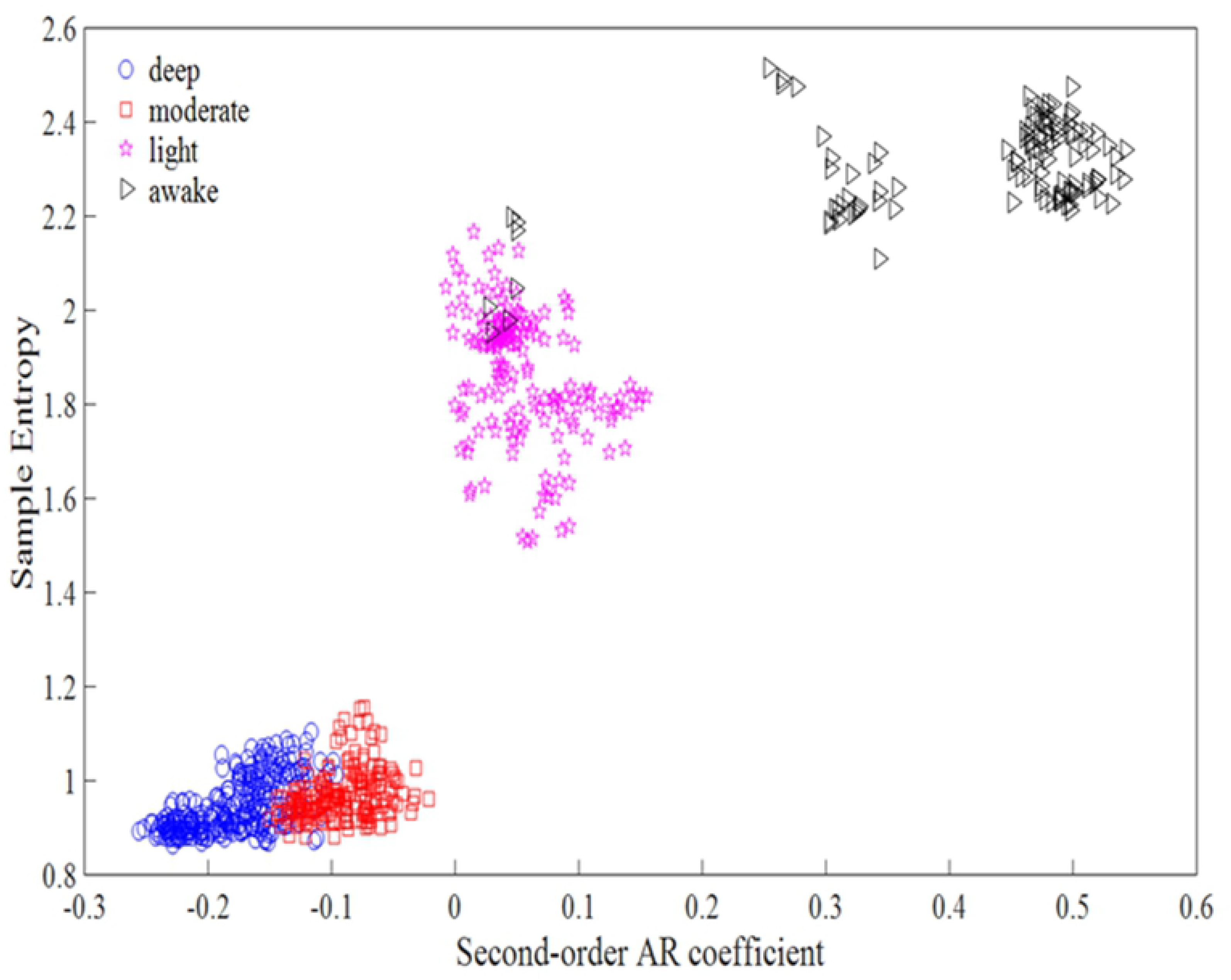
Distribution of sample entropy and second-order AR coefficient. The data was obtained from EEG of one patient, the x-coordinate is the value second-order AR coefficient and y-coordinate is the value of SampEn.

We compare the classification effect of WKNN with KNN and SVM. As shown in Table 2, whether it is a class of feature vectors or two types of feature vectors, the accuracy of WKNN is higher than the accuracy of SVM and KNN. The results show that WKNN is suitable for the classification requirements of this study and can achieve accurate results.

**Table.2.** WKNN and KNN, SVM classification results

| method | sampEn | ArIndex | sampEn+ArIndex(Acc) |
| --- | --- | --- | --- |
| SVM | 77.10% | 72.97% | 79.61% |
| KNN | 76.65% | 84.12% | 87.35% |
| WKNN | 78.13% | 86.25% | 88.62% |

The sensitivity and specificity of the WKNN classifier are obtained by calculation. As can be seen from the results shown in Table 3, the sensitivity and specificity of the WKNN classifier under different anesthesia states are above 0.8. WKNN has very high sensitivity and specificity, indicating that it can obtain accurate classification results.

**Table.3.**
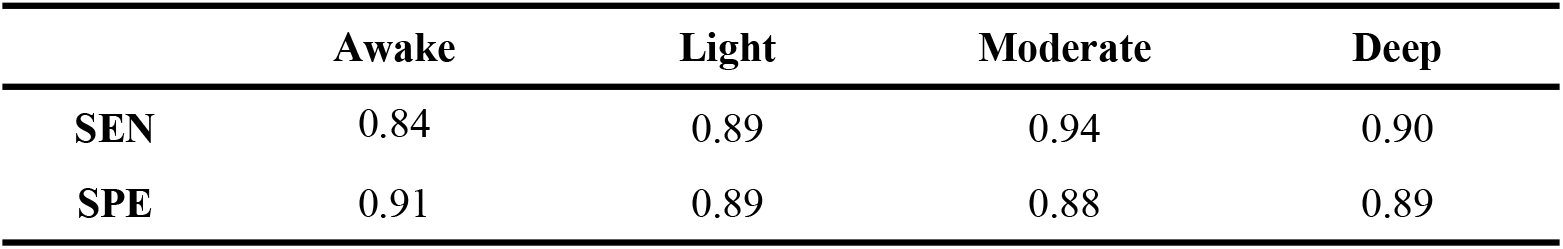
WKNN classifier sensitivity and specificity under different anesthesia states

|  | Awake | Light | Moderate | Deep |
| --- | --- | --- | --- | --- |
| **SEN** | 0.84 | 0.89 | 0.94 | 0.90 |
| **SPE** | 0.91 | 0.89 | 0.88 | 0.89 |

### Results analyzing

Sample entropy as a feature of the response EEG signals complexity has been widely used in EEG analysis [36,37]. Through analyzing the SampEn of the EEG signals found that the correlation coefficient between SampEn of the EEG signals and BIS value is as high as 0.8. Fig.3a shown that the trend of sample entropy and BIS values. It can be seen from the results that sample entropy can accurately reflect the depth of anesthesia. AR model can well represent the internal features and information of the signal. Recently, AR model has been widely used in EEG processing. We choose the optimal AR coefficient, which reduces the complexity of the redundancy and the calculation of the features. The recognition rate of different AR coefficient combinations is shown in the Fig.5b. The WKNN method is especially suitable for the multi-classification problem of sample non-uniformity, and it is very important to choose the favorable k value. Taking the different K values into account, we calculated the recognition rate corresponding to different K values, and obtained the recognition rate results as shown in Fig.5a. Through the relationship between the K values and the corresponding recognition rate in Fig.5a, the K value is selected as seven. Finally, through the AR coefficient and SampEn as feature vectors, the WKNN classifier is used to classify the depth of anesthesia into four states. It can be seen from the recognition rate (Fig.6) that the method can be used for the monitoring of DOA.

According to the research method in this paper, we calculate the SampEn of EEG signals at different anesthesia depths. From Fig.3a, we can clearly see that the SampEn of the EEG signals gradually decreases when the depth of anesthesia increases. This phenomenon indicates that the activity level of the brain decreases with the increase of the depth of anesthesia, which is consistent with the working state of the brain. Through extracting AR coefficients and the correlation between AR coefficients and BIS values, as shown in Table.1, it is found that the first five order AR coefficients are correlated with BIS. Through condition optimization, two to five order AR coefficients were selected as feature vectors. Further analysis of sample entropy and second-order AR coefficients to obtain their box and scatter plot, as shown in Fig.3b and Fig.4b, both characteristics can well complete the classification of depth of anesthesia. The sample entropy and the second-order AR coefficient of five patients who under different anesthesia conditions were calculated separately. It can be seen from Table.4 and Table.5 that the sample entropy and the second-order AR coefficient have significant differences under different depth of anesthesia.

**Table.4.** Sample entropy under different DOA

| Patients | sample entropy |  |  |  |
| --- | --- | --- | --- | --- |
|  | Awake | moderate | light | deep |
| 1 | 2.36($\pm 0.16$) | 1.87($\pm 0.15$) | 1.01($\pm 0.11$) | 0.91($\pm 0.05$) |
| 2 | 1.86($\pm 0.11$) | 1.31($\pm 0.13$) | 1.01($\pm 0.07$) | 0.98($\pm 0.08$) |
| 3 | 1.81($\pm 0.15$) | 1.61($\pm 0.19$) | 1.07($\pm 0.12$) | 0.91($\pm 0.09$) |
| 4 | 2.12($\pm 0.10$) | 1.39($\pm 0.12$) | 1.09($\pm 0.08$) | 1.03($\pm 0.04$) |
| 5 | 2.52($\pm 0.09$) | 1.51($\pm 0.11$) | 1.32($\pm 0.18$) | 1.19($\pm 0.06$) |

**Table.5.** Second order AR coefficient under different DOA

| Patients | second order AR coefficient |  |  |  |
| --- | --- | --- | --- | --- |
|  | Awake | moderate | light | deep |
| 1 | 0.48($\pm 0.09$) | 0.09($\pm 0.08$) | -0.09($\pm 0.06$) | -0.21($\pm 0.06$) |
| 2 | 0.69($\pm 0.05$) | 0.24($\pm 0.09$) | -0.04($\pm 0.09$) | -0.08($\pm 0.07$) |
| 3 | 0.51($\pm 0.23$) | 0.42($\pm 0.18$) | 0.07($\pm 0.08$) | -0.07($\pm 0.08$) |
| 4 | 0.46($\pm 0.26$) | 0.11($\pm 0.06$) | -0.03($\pm 0.04$) | -0.07($\pm 0.03$) |
| 5 | 0.87($\pm 0.14$) | 0.19($\pm 0.08$) | -0.09($\pm 0.08$) | -0.09($\pm 0.04$) |

Finally, we select k=7, and use the WKNN classifier to classify the DOA. In the classification process, the ratio of training samples to test samples is 4:1. We can see the classification results in Fig.6, the method can well complete the classification of DOA. Through analyzing the classification results, the distribution of misjudged samples under four anesthesia conditions is obtained. From Fig.7, it can be found that a large part of the awake state is misjudged as light anesthesia. The reason for this phenomenon may be that the noise of EEG signals in awake state is too much to be filtered thoroughly, which leads to the decrease of sample entropy [38].

Using the selected eigenvalues to compare the three classifiers commonly used in WKNN, KNN and SVM, as shown in Table.3, WKNN can achieve better classification results when the sample size distribution is not uniform. Through calculating the sensitivity and specificity of WKNN classifier, it is found that WKNN has good sensitivity and specificity. The above results show that WKNN can well divide the depth of anesthesia into four states: awake, mild, moderate and deep anesthesia.

## Discussion

In this study, we propose a new method for monitoring the DOA, which based on SampEn and AR coefficients as feature vectors combined with the WKNN classifier. Firstly, the original EEG signal is segmented by sliding window, the window length is 2560 (10 seconds of EEG signals) and step size is 512 (2 seconds of EEG signals). Then, the sample entropy and AR coefficients of EEG signals are calculated, and the optimal AR coefficients are selected. The characteristics reflecting the depth of anesthesia were extracted successfully and the highest correlation index between these characteristics and BIS index is over 0.8. Finally, the optimal K value is selected and the depth of anesthesia is successfully divided into four states: awake, mild, moderate and deep anesthesia by WKNN classifier. The classification accuracy of this method is over 88%. To sum up, the combination of sample entropy and AR coefficient can monitor the depth of anesthesia accurately and effectively.

## Author Contributions

conceptualization, X.L.; methodology, X.L. and F.Z.; software, F.Z; validation, X.L.; formal analysis, F.Z. and C.H.; investigation, X.W. and X.L; resources, D.H.; data curation, F.Z. and P.D.; writing—original draft preparation, F.Z.; writing—review and editing, X.L. and F.Z.; visualization, F.Z.; supervision, X.L.; project administration, X.L.; funding acquisition, X.L”.

## Acknowledgments

The authors gratefully acknowledge the help of all the volunteers in this study. We also would like to thank department help in EEG data collection.

